# HCMV miR-US22 down-regulation of EGR-1 regulates CD34+ hematopoietic progenitor cell proliferation and viral reactivation

**DOI:** 10.1101/645374

**Authors:** Iliyana Mikell, Lindsey B. Crawford, Meaghan H. Hancock, Jennifer Mitchell, Jason Buehler, Felicia Goodrum, Jay A. Nelson

## Abstract

Reactivation of latent Human Cytomegalovirus (HCMV) in CD34+ hematopoietic progenitor cells (HPCs) is closely linked to hematopoiesis. Viral latency requires maintenance of the progenitor cell quiescence, while reactivation initiates following mobilization of HPCs to the periphery and differentiation into CD14+ macrophages. Early growth response gene 1 (EGR-1) is a transcription factor activated by Epidermal growth factor receptor (EGFR) signaling that is essential for the maintenance of CD34+ HPC self-renewal in the bone marrow niche. Down-regulation of EGR-1 results in mobilization and differentiation of CD34+ HPC from the bone marrow to the periphery. In the current study we demonstrate that the transcription factor EGR-1 is directly targeted for down-regulation by HCMV miR-US22 that results in decreased proliferation of CD34+ HPCs and a decrease in total hematopoietic colony formation. We also show that an HCMV miR-US22 mutant fails to reactivate in CD34+ HPCs, indicating that expression of EGR-1 inhibits viral reactivation during latency. Since EGR-1 promotes CD34+ HPC self-renewal in the bone marrow niche, HCMV miR-US22 down-regulation of EGR-1 is a necessary step to block HPC self-renewal and proliferation to induce a cellular differentiation pathway necessary to promote reactivation of virus.

**Author summary:** Human cytomegalovirus (HCMV) is a widespread herpesvirus that persists in the host and remains a significant cause of morbidity and mortality in solid organ and stem cell transplant patients. HCMV latency is complex, and the molecular mechanisms for establishment, maintenance, and reactivation from latency are poorly understood.

Quiescent stem cells in the bone marrow represent a critical reservoir of latent HCMV, and the mobilization and differentiation of these cells is closely linked to viral reactivation from latency. HCMV encodes small regulatory RNAs, called miRNAs that play important roles in the regulation of viral and cellular gene expression. In this study, we show that HCMV miR-US22 targets Early growth response gene 1 (EGR-1) a host transcription factor that is necessary for stem cell quiescence and self-renewal in the bone marrow. Expression of this miR-US22 down-regulates expression of EGR-1 that reduces CD34+ HPCs proliferation and total hematopoietic colony formation. An HCMV miR-US22 mutant is unable to reactivate from latency suggesting that the ability of the miRNA to disrupt CD34+ HPC renewal in the bone marrow niche to initiate a differentiation pathway is critical for viral reactivation.

## Introduction

Human cytomegalovirus (HCMV) remains a significant cause of morbidity and mortality in solid organ and hematopoietic stem cell transplant patients [1–3]. CD34+ hematopoietic progenitor cells (HPCs) represent a critical reservoir of latent HCMV in the transplant recipient, providing a source of virus for dissemination to visceral organs. HCMV latency is complex, and the mechanisms for establishment and maintenance of HCMV latency and reactivation of virus are poorly understood at the molecular level. HCMV reactivation is exquisitely linked to CD34+ HPC hematopoiesis and differentiation into myeloid lineage cells [4, 5]. Viral regulation of the CD34+ HPC hematopoiesis program is considered a major determinant of HCMV latency and reactivation. Activation of growth factor receptor signaling that induces transcriptional reprogramming is necessary to both maintain CD34+ HPCs in a quiescent state and induce myelopoiesis. Viral regulation of these events determines whether the HCMV remains latent or initiates the reactivation program.

Establishment of latency likely involves both the expression of viral factors suppressive of replication and a cellular environment that supports the epigenetic silencing of the viral genome (reviewed in [6, 7]). The latent state is characterized by the absence of the gene expression repertoire that is otherwise associated with virion production in fibroblasts [8]. Reactivation of viral gene expression is closely tied to mobilization of HPCs to the periphery and differentiation into CD14+ monocytes [9–11]. In infected individuals the viral genome is maintained at very low copy numbers, and detection of viral gene expression *in vivo* is challenging, hence experimental models of cultured CD34+ HPCs have been instrumental in studying molecular models of latency and reactivation (discussed in [12]).

Early growth response gene 1 (EGR-1) is a member of a family of sequence-specific zinc finger transcription factors that was originally characterized as an oncogene [13–16] but was later observed to be important in multiple cellular processes, including cell proliferation, differentiation, and apoptosis (reviewed in [17]). EGR-1 is activated by epidermal growth factor receptor (EGFR) signaling that is an important regulator of normal hematopoiesis through the control of key cell cycle regulators, cytokines, and co-stimulatory molecules [18, 19]. EGR-1 expression in CD34^+^ HPCs promotes “stemness” (self-renewal and lack of differentiation) in the bone marrow niche [18]. Consequently, deletion of the EGR-1 gene in mice promotes CD34+ HPC differentiation and migration to the periphery [18]. Importantly, EGR-1 plays a dual role in the development of myeloid cells during hematopoiesis. In a subset of progenitor cells, expression of Egr-1 inhibits the differentiation of myeloid precursor cells along the macrophage lineage [20], while in monocytes EGR-1 potentiates terminal macrophage differentiation [21]. Therefore, the timing of EGR-1 expression is an important determinant of CD34+ HPC myelopoiesis.

EGFR and downstream PI3K signaling are important for establishing and maintaining a latent infection in CD34+ HPCs [22, 23]. HCMV stimulates EGFR upon entry into CD34+ HPCs and then is thought to induce an environment primed for the establishment of latency. Contrary to infection of fibroblasts that support virus replication, EGFR cell surface levels are transiently increased during infection of CD34+ cells [22]. The HCMV proteins UL138 and UL135 oppose one another in regulating the trafficking of EGFR and, thus, its capacity for signaling. UL135 targets EGFR for turnover through its interaction with the host adapter proteins Abi-1 and CIN85 [24]. These UL135-host protein interactions and the attenuation of EGFR and its downstream signaling are important for HCMV reactivation in CD34+ HPCs [22, 24].

HCMV, similar to other herpesviruses, encodes multiple miRNAs [25, 26] expressed during both the viral lytic and latent phases of infection (reviewed in [27]). HCMV miRNAs regulate the expression of cellular and viral genes involved in viral replication [28, 29], formation of the viral assembly compartment [30], inhibition of proinflammatory cytokines production and release [31, 32], immune evasion [33–35], and promotion of cell survival [36]. A common theme among herpesviruses is that viral miRNAs target both viral and cellular transcripts in order to restrict acute viral replication and maintain the latent state. For example, HCMV miR-UL112-1-3p directly targets the 3’ UTR of IE72 that limits lytic gene expression and tempers virus replication [28]. Conversely, both HSV and EBV induce cellular miRNAs to target viral lytic switches of HSV-1 (ICP0) directly [37], or EBV (BZLF1) indirectly [38]. Regulation of the cell cycle and the cellular differentiation state are also a critical determinant of whether HCMV maintains a latent state in CD34+ HPCs or stimulates cellular differentiation resulting in viral reactivation. HCMV miR-US25-1 was shown to target five cell cycle genes, including cyclin E2, and several genes that modify DNA chromatin [39]. Lastly, HCMV miR-US25-2-3p was shown to decrease both viral and cellular DNA synthesis and cell proliferation [40, 41]. These data suggest that HCMV miRNAs promote a cellular state associated with reduced replication in order to maintain viral latency. Thus, the regulation of both viral and cellular genes by HCMV miRNAs provides an important mechanism to maintain latency or initiate reactivation without the expression of viral proteins that could trigger an immune response to the infected cell.

In the current study, we demonstrate that HCMV miR-US22 efficiently down-regulates EGR-1 expression that results in a decrease in total hematopoietic colony formation and progenitor cell proliferation. Additionally, mutation of miR-US22 in the virus significantly reduced the ability of HCMV to reactivate in CD34+ HPCs. These data indicate that miR-US22 regulation of EGR-1 expression is a necessary step in viral reactivation from latency.

## Results

### HCMV miR-US22 Down-Regulates Expression of EGR-1

Bioinformatics analysis of potential HCMV miRNA cellular targets indicates that EGFR signaling is one of the most heavily targeted signaling pathways (data not shown). Since EGR-1 is activated downstream of EGFR signaling and is a key transcription factor regulating stemness of CD34+ HPCs in the bone marrow niche, we examined whether HCMV miRNAs functionally target EGR-1 activity. For this experiment, several HCMV miRNA mimics were co-transfected with an EGR-1 luciferase reporter into HEK293 cells in the presence or absence of EGF to examine their effect on promoter activation. Several of the HCMV miRNAs significantly altered luciferase expression from the EGR-1 reporter (Fig 1). A 3-4 fold decrease in luciferase reporter activity was observed with transfection of miR-US22. In contrast, transfection of miR-US5-1 and miR-UL112-3p resulted in up to a two-fold increase in promoter activity. We focused on miR-US22-mediated EGR-1 downregulation since miR-US5-1 and miR-UL112-3p upregulation of EGR-1 activity is the focus of another study. Analysis of the 3’ UTR of the EGR-1 mRNA revealed a miR-US22 seed sequence binding site (Fig 2A). In order to validate the miR-US22 target sequence, two base pairs were mutated in the EGR-1 3’ UTR cloned in a luciferase reporter plasmid (Fig 2A). Co-transfection of HEK293 cells with WT and mutant EGR-1 luciferase constructs and miR-US22 mimic indicated that mutation of the miR-US22 target sequence fully restored EGR-1 reporter activity, validating this sequence as a miR-US22 target site (Fig 2B). To determine if miR-US22 can reduce EGR-1 protein expression in cells, HEK293 cells and normal human dermal fibroblasts (NHDF) were transfected with a miR-US22 mimic or EGR-1 siRNA, followed by serum starvation and addition of EGF to induce EGR-1 expression. Both the miR-US22 mimic and EGR-1 siRNA reduced EGR-1 protein levels in both HEK293 and NHDF cells by 10-fold and 3-fold, respectively (Fig 2C).

**Fig 1.**
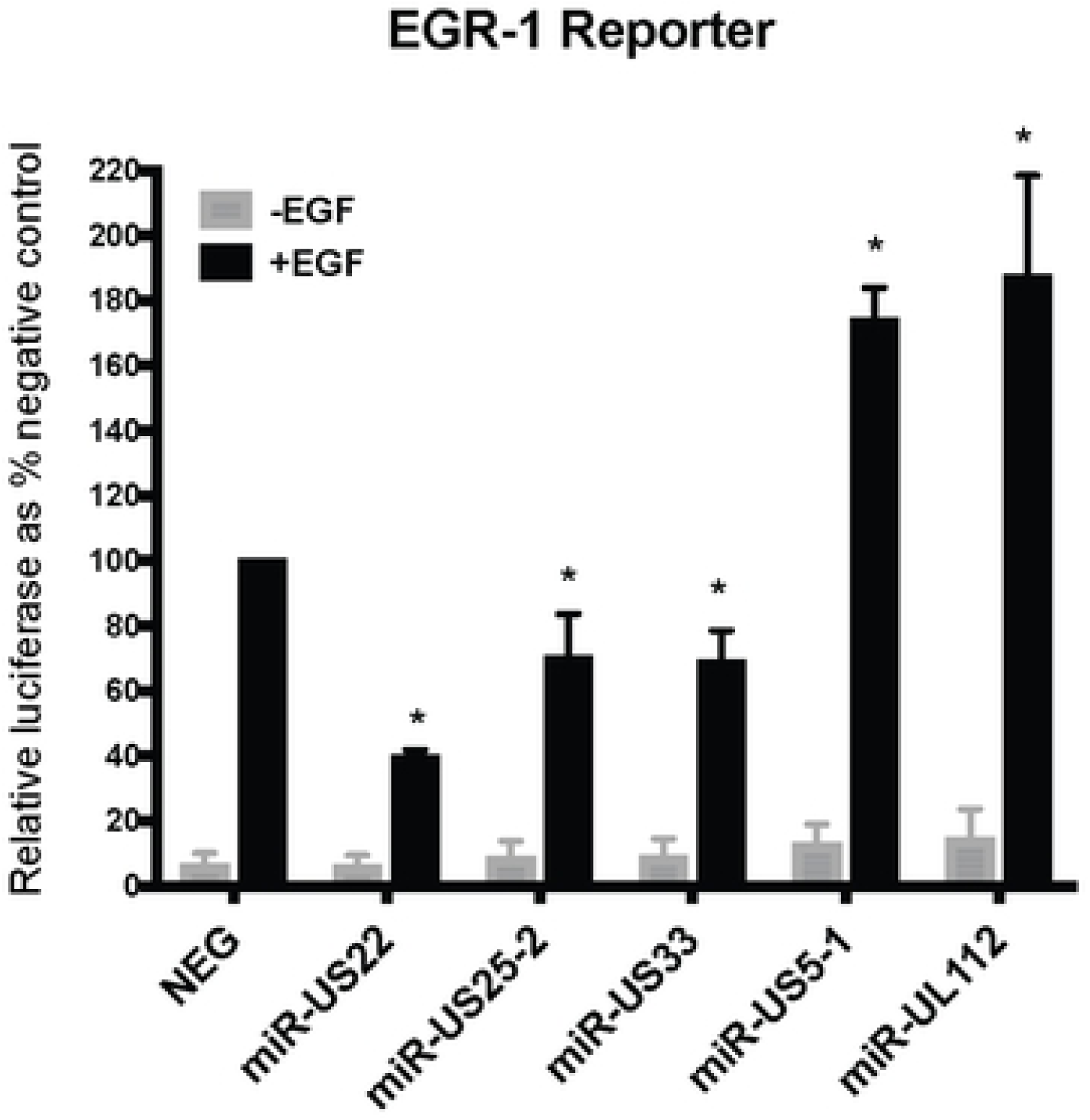
HCMV miRNAs affect EGF-mediated signaling to EGR-1. An EGR-1 luciferase reporter construct was transfected in HEK293 cells along with negative siRNA control (NEG) or HCMV miRNA mimics. After 24 hours, the cells were serum-starved for 4 hours followed by 4 hours of EGF treatment (5 ng/mL). Cells were then harvested and luciferase expression was assessed. Data from 3 independent experiments are graphed as mean ± SD; * p<0.01 compared to NEG – transfected cells (unpaired t test).

**Fig 2.**
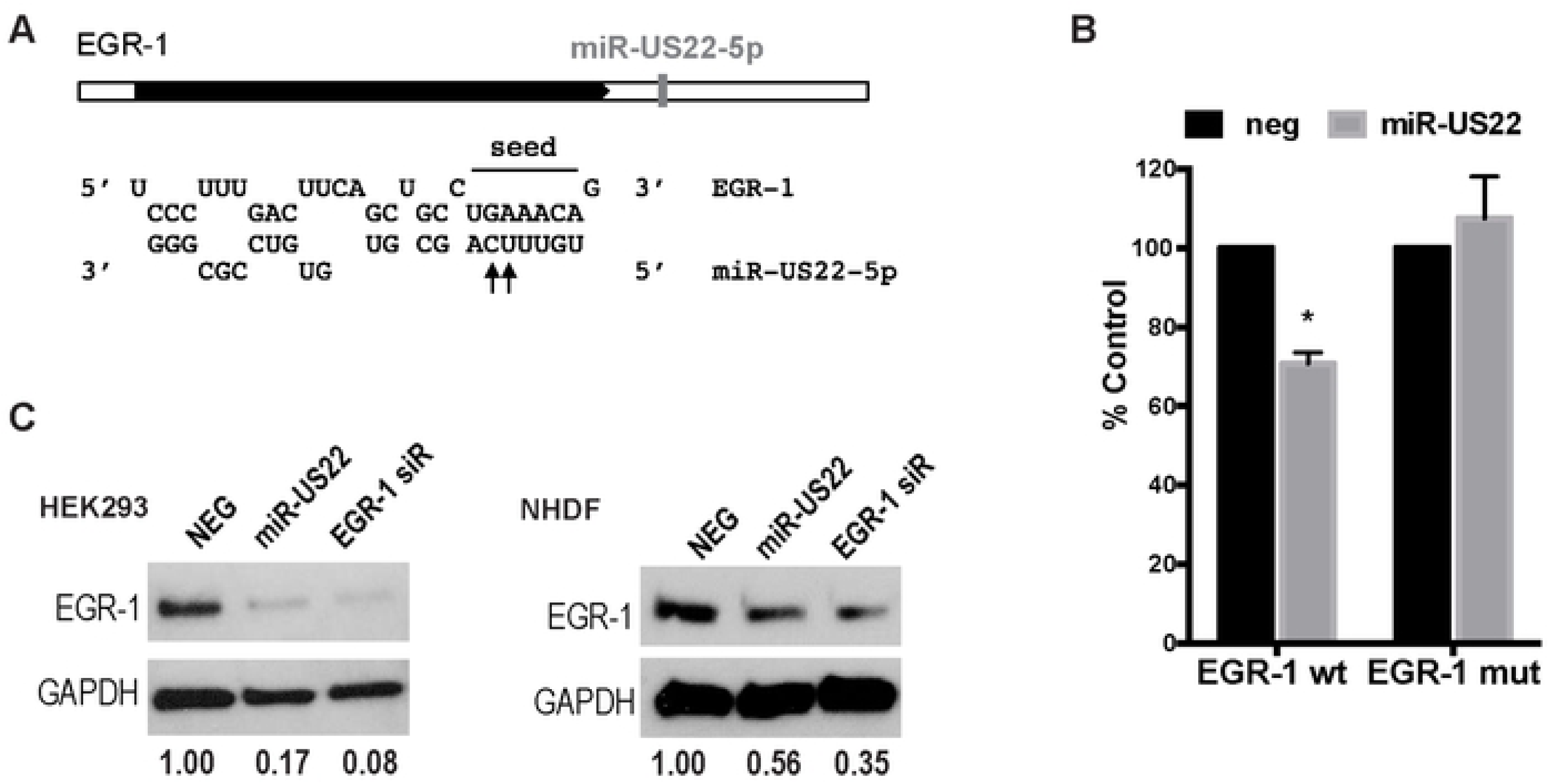
EGR-1 3’UTR is targeted by HCMV miR-US22. (A) One miR-US22 target site is present in the EGR-1 3’UTR. The seed sequence is indicated; the arrows label the bases that were mutated to disrupt miR-US22 binding to the target site. (B) The miR-US22 target site is required for EGR-1 down-regulation. Dual luciferase reporter containing EGR-1 3’UTR with wild type target site or mutated target site were co-transfected with the indicated mimics, and assessed for luciferase expression 24 hours later. Data graphed as mean ± SD from 3 independent experiments; * p=0.005 compared to NEG - transfected cells (unpaired t test). (C) HEK293 or NHDF cells were transfected with NEG siRNA control, miR-US22 mimic, or EGR-1 siRNA for 48 hours, after which the cells were serum starved (4 hours), treated with 50ng/mL EGF (1 hour), and harvested for immunoblot (IB) analysis. Protein concentrations were normalized to GAPDH, and relative levels are displayed.

In order to validate miR-US22 targeting of EGR-1 during infection, a recombinant HCMV construct was designed with mutations in miR-US22 that disrupt the miRNA expression. To construct the HCMV ΔmiR-US22, BAC recombineering was used to introduce 5 silent mutations that do not disrupt the US22 ORF (Fig 3A) into the stem loop of the miRNA in HCMV TB40E (Fig 3B). We have previously published that disruption of the miRNA stem loop inactivates the function of other HCMV miRNAs [30]. Sequence analysis of HCMV ΔmiR-US22 indicated that the only difference between the mutant and WT TB40E virus was the 5 bases introduced into the miR-US22 stem loop (data not shown). The HCMV TB40E ΔmiR-US22 lacked expression of miR-US22 (Fig 3C), exhibited WT growth in NHDF cells (Fig 3D), and did not impair US22 expression (Fig 3E). In order to determine the effect of the miR-US22 mutation on EGR-1 expression, western blot analysis of EGR-1 was performed on HCMV WT and ΔmiR-US22 TB40E infected NHDF or aortic endothelial cells (AEC) at 2, 3, 4 and 6 days post infection (dpi). Infection with the HCMV ΔmiR-US22 resulted in 2-3-fold increase in EGR-1 protein expression at all time points compared to WT infection in both NHDFs (Fig 4A) and AECs (Fig 4B). Additionally, transfection of exogenous miR-US22 mimic in cells infected with HCMV ΔmiR-US22 reversed the effect of the mutation and resulted in markedly reduced EGR-1 protein levels, compared to cells transfected with a negative control (Fig 4C). The above data indicate that EGR-1 is a miR-US22 target during HCMV infection.

**Fig 3.**
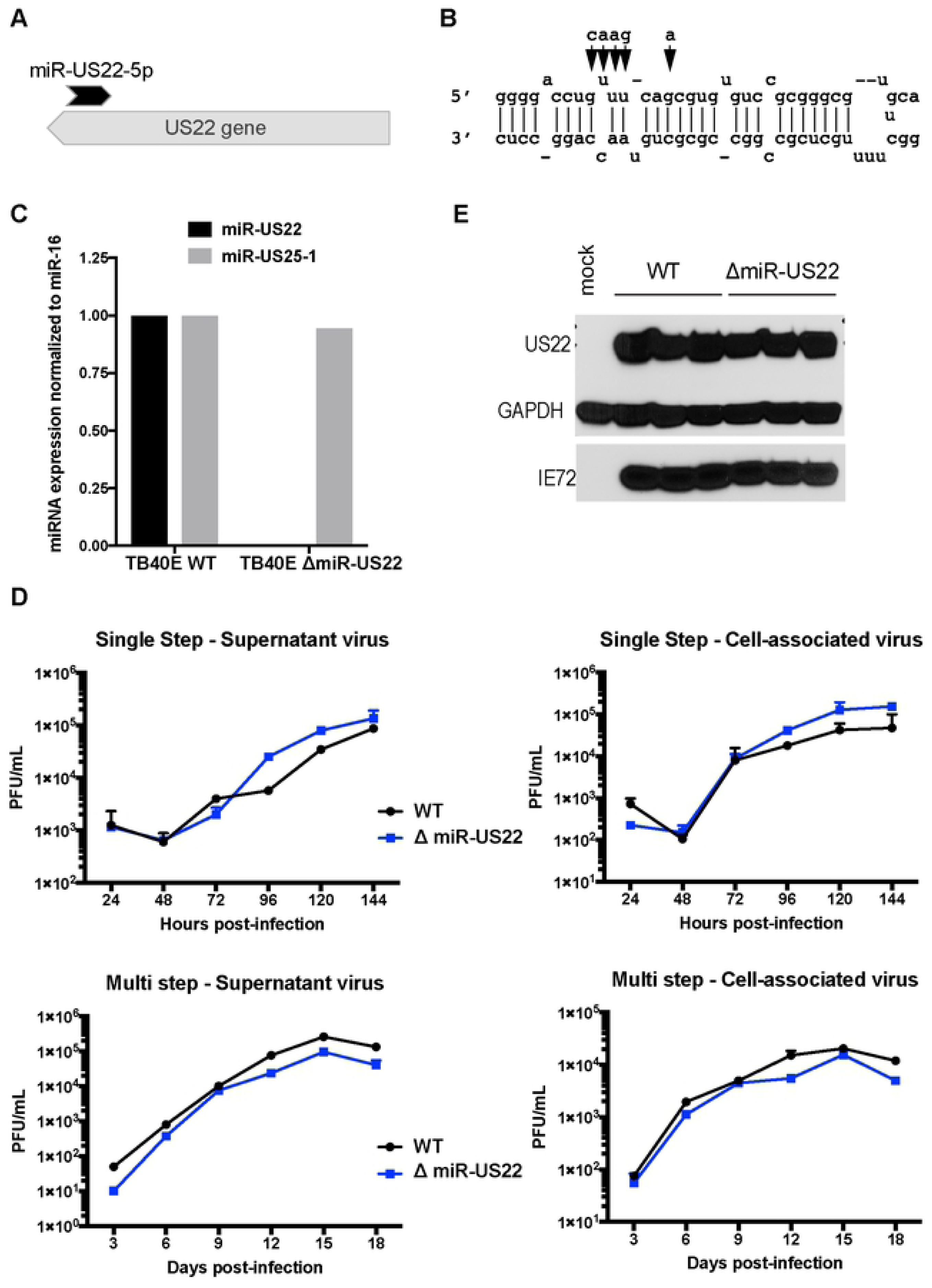
Characterization of TB40E miR-US22 mutant virus. (A) Relative location of miR-US22 within the US22 gene. (B) Introduction of point mutations (arrows) in TB40E to disrupt miR-US22 hairpin formation. (C) Loss of miR-US22 expression in NHDFs infected with the TB40E ΔmiR-US22 virus. NHDFs were infected with either WT TB40E or ΔmiR-US22 virus (MOI=3), and miRNA levels were determined 4 days post infection by stem-loop specific RT-PCR. miR-US25-1 expression is shown as a control. (D) Mutating miR-US22 in TB40E has no effect on viral growth in fibroblasts. NHDF were infected in duplicate with either WT TB40 (black circles) or TB40E ΔmiR-US22 (blue squares) at MOI=3 for single step or MOI=0.05 for multi step. Plaque forming units (pfu) / mL were quantified from samples collected at the indicated time points for supernatant virus and cell-associated virus (E) The point mutations in miR-US22 do not affect US22 protein expression. NHDFs were infected with either WT TB40E or ΔmiR-US22 TB40E, and protein lysates were collected 4 days later for IB analysis. 3 replicates are shown.

**Fig 4.**
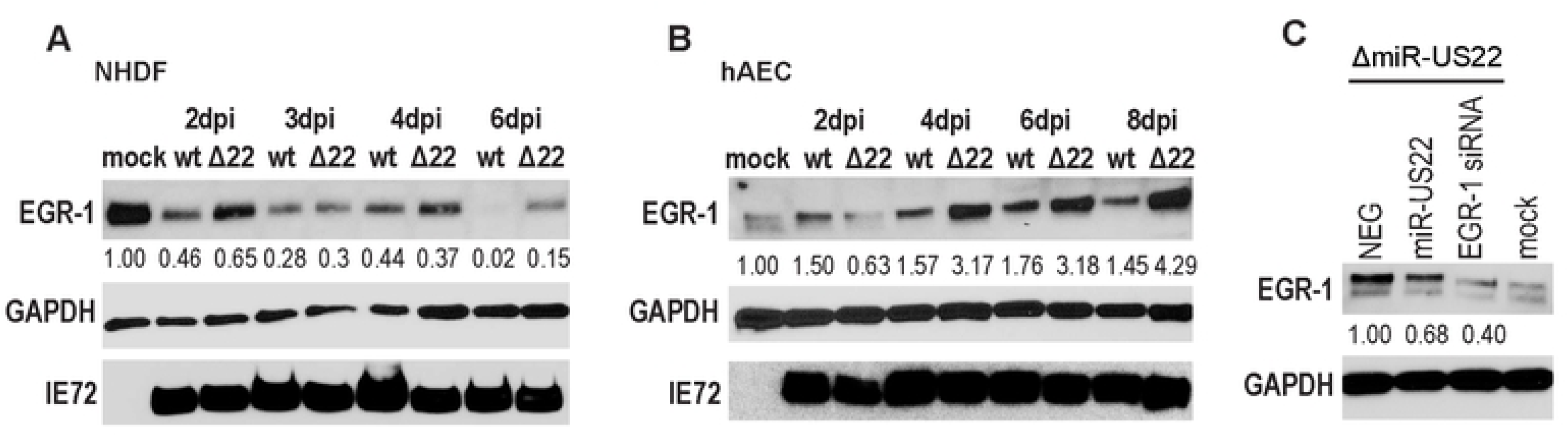
miR-US22 targets endogenous EGR-1 during HCMV infection. (A, B) TB40E miR-US22 mutant virus fails to down-regulate endogenous EGR-1 levels during infection. NHDF (A) or AEC (B) were infected with TB40E WT or ΔmiR-US22 virus (MOI = 3). Cell lysates were harvested at the indicated timepoints for IB analysis (C) miR-US22 mimic downregulates EGR-1 in cells infected with the miR-US22 mutant virus. AECs were transfected with the indicated siRNA or miRNA mimics. After 24 hours, the cells were infected with TB40E ΔmiR-US22, and harvested 4 days later for IB analysis. Protein levels were normalized to GAPDH, and relative levels are displayed.

### HCMV miR-US22 reduces CD34+ HPC proliferation

Since HCMV latent infection of CD34+ HPCs results in decreased myeloid colony formation, and expression of EGR-1 is a major determinant in the proliferative capacity of progenitor cells, we examined the direct effects of miR-US22 on myelopoiesis. CD34+ HPCs were transfected with either a GFP-containing plasmid that expresses miR-US22, an shRNA to EGR-1, or a negative control, followed by sorting 24 hours later for GFP+ cells. The sorted cells were placed in myeloid colony formation support medium and analyzed by microscopy at 7 days post plating (Fig 5A). Transfection of miR-US22 or shRNA to EGR-1 significantly reduced total myeloid colony formation by 30% and 36%, respectively. Analysis of the types of myeloid colonies that were negatively affected by miR-US22 down-regulation of EGR-1 indicated a decrease in both CFU-GM and BFU-E colonies (Fig 5B). The ratio of total myeloid to erythroid colonies was unchanged, suggesting that the effect of the miRNA was on progenitor cell proliferation rather than altering HPC differentiation or lineage commitment. In order to determine whether miR-US22 altered the ability of CD34+ HPCs to proliferate, cells transfected with a miR-US22 expression plasmid, shRNA to EGR-1, or a negative control were sorted and plated in stem cell cytokine-enriched media, followed by quantitation of cell numbers at 3- and 7-days post transfection (Fig 5C). Transfection with either miR-US22 or shRNA to EGR-1 reduced CD34+ HPC proliferation by 2 and 4-fold respectively in comparison to a control plasmid or mock treated cells. These data indicate that miR-US22 and an shRNA to EGR-1 significantly reduces the proliferation of CD34+ HPCs.

**Fig 5.**
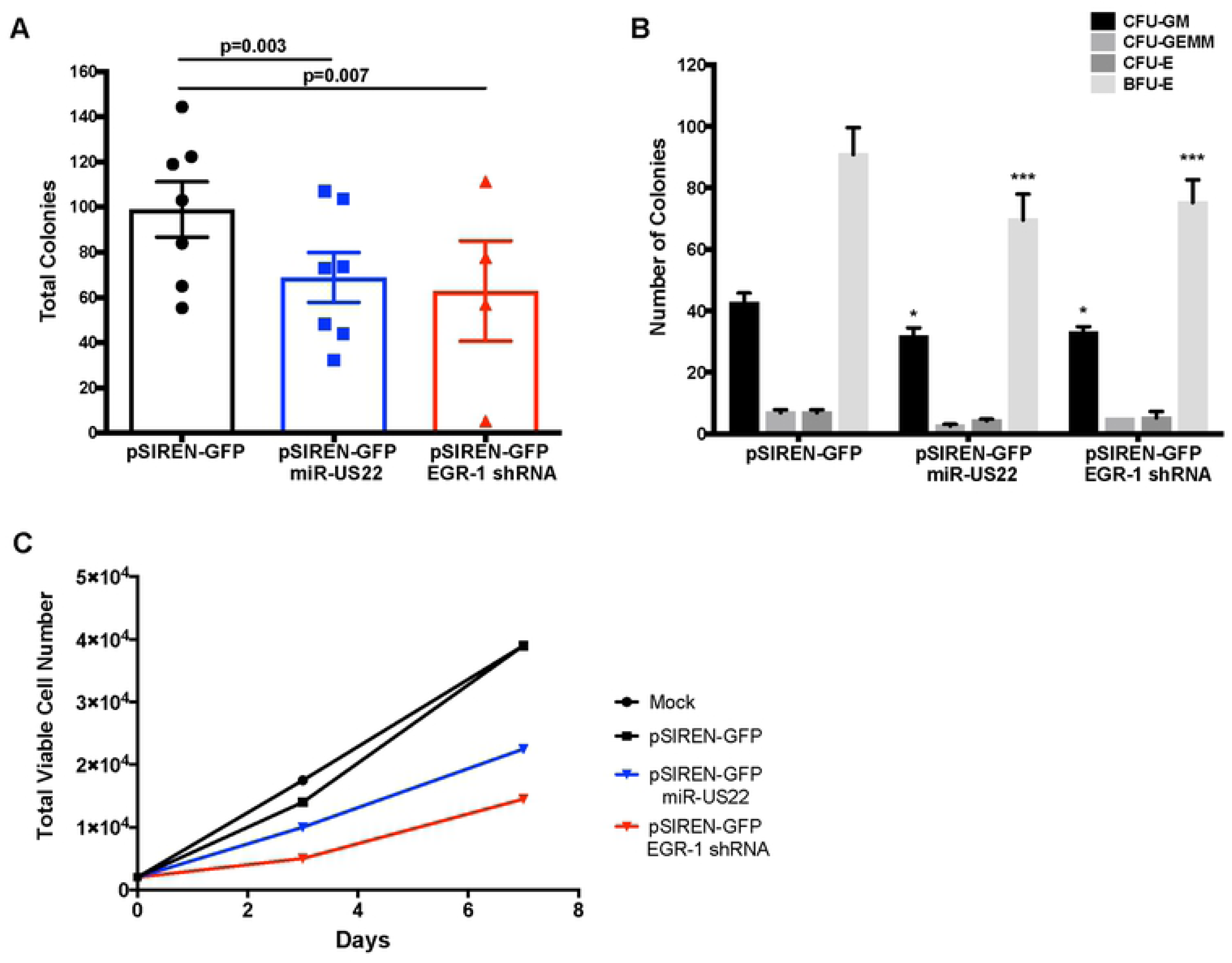
miR-US22 down-regulation of EGR-1 reduces CD34+ HPC proliferation. CD34+ HPCs were transfected with pSIREN-GFP, pSIREN-GFP-miR-US22, or pSIREN-GFP-EGR-1shRNA for 24 hours using Amaxa. Viable CD34+ GFP+ HPCs were isolated by FACS and analyzed for proliferation and differentiation. (A, B) Isolated HPCs were plated on Methocult H4434 (Stem Cell Technologies) at 500 cells/mL in triplicate, and counted at 7 and 14 days. Data shown are the total number of colonies at 7 days (A) for n=7 (miR-US22) or n=3 (EGR-1 shRNA) independent experiments, or separate myeloid (CFU-GM, CFU-GEMM) and erythroid (CFU-E and BFU-E) colonies at 14 days (B). Significance was calculated using paired t-test; * p=0.019, *** p<0.001 compared to pSIREN-GFP control transfected cells. (C) Isolated CD34+ HPCs were plated in SFEMII supplement with hematopoietic cytokines, and counted at day 3 and day 7. Total viable cell number is shown.

### HCMV miR-US22 is required for viral reactivation in latently infected CD34+ HPCs

Since herpesvirus miRNAs play key roles in latency and reactivation, HCMV miRNAs expressed at high levels during lytic infection were examined for expression in CD34+ cells. HCMV TB40E-GFP infected CD34+ HPCs were sorted for GFP and incubated for 14-days post infection (dpi) on stromal cell support followed by extraction of RNA. Analysis of miRNA expression in HCMV latently infected CD34+ HPCs indicated that several viral miRNAs are detected with varying levels of expression. The latently expressed miRNAs include miRs -UL22A, -UL112-3p, -UL148D, -US5-1, -US5-2, -US25-1, and -US25-2-3p (Fig 6). HCMV miRs -UL22A, -UL112-3p and -US25-1 represented some of the most abundant miRNAs. In contrast, miRs -UL36, -US4, -US22, -US29, and -US33 were not detected in latently infected cells (data not shown). Therefore, although miR-US22 is expressed during acute infection, the HCMV miRNA is not expressed in latently infected CD34+ HPCs.

**Fig 6.**
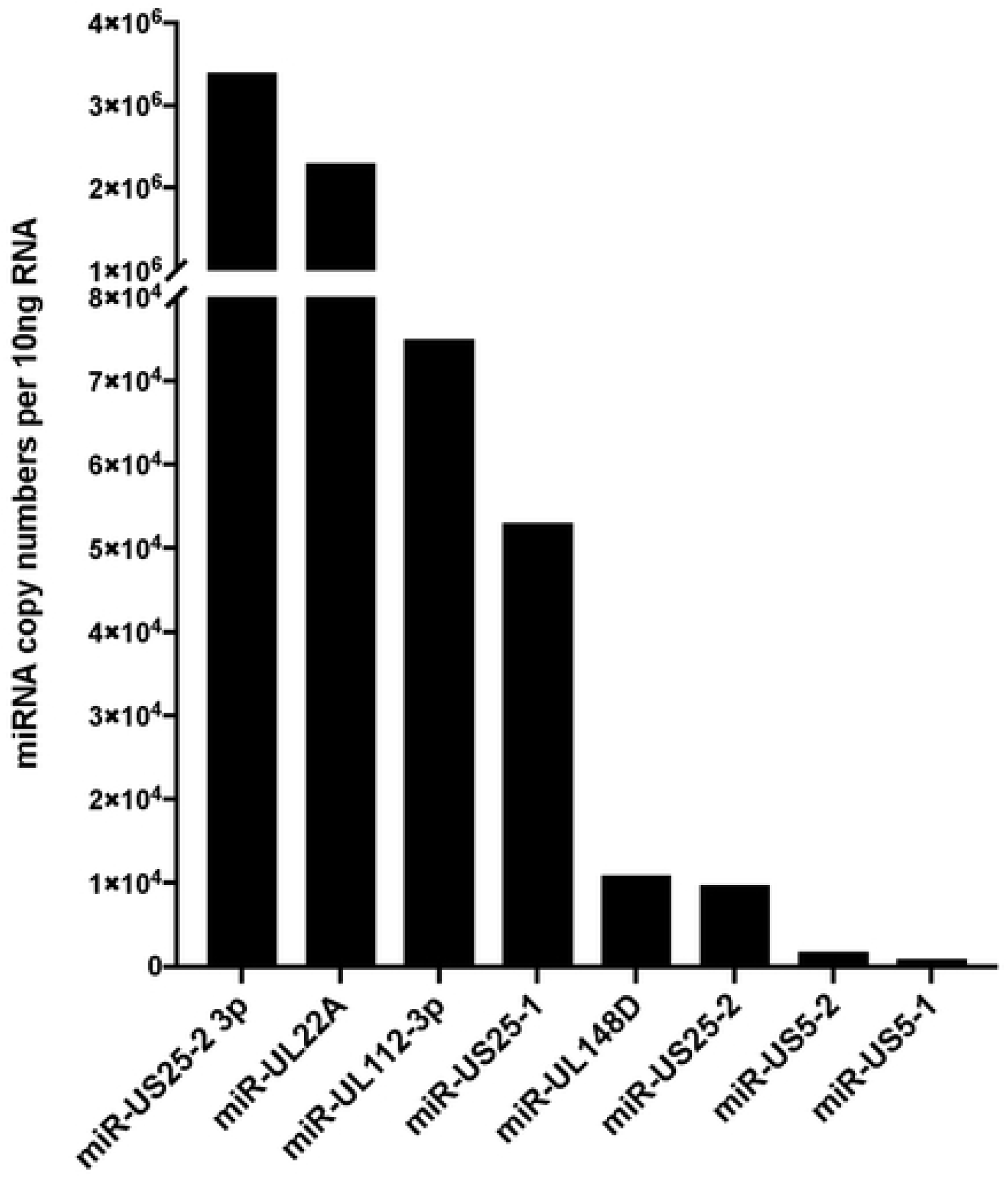
Expression of HCMV-encoded miRNAs in latently infected CD34+ HPCs. CD34+ HPCs were infected with WT HCMV TB40E for 48 hours, and then FACS-isolated for viable CD34+ GFP+ HPCs. Sorted cells were plated on stromal cell support for 12 additional days to establish HCMV latency, and HCMV miRNA levels were detected in 10ng RNA from infected cells by stem-loop RT-PCR for viral miRNAs.

Since EGR-1 plays a critical role in the maintenance of progenitor cell stemness, and HCMV latency and reactivation is integrally linked to cellular differentiation, the role of miR-US22 in viral latency and reactivation was examined in CD34+ HPCs. To determine whether miR-US22 is required for the reactivation process, CD34^+^ HPCs were infected with either HCMV WT TB40E-GFP or the ΔmiR-US22 mutant, and were sorted for GFP expression to acquire a pure population of infected cells. Infected CD34+ HPCs were seeded into long-term bone marrow cultures using a stromal cell support shown to maintain stem cells. After 12 days in culture, the cultures were split. Live cells from half of the culture were seeded by limiting dilution onto monolayers of fibroblasts in cytokine-rich media to promote myeloid differentiation. The frequency of infectious centers, determined by extreme limiting dilution assay, was calculated from the fraction of GFP+ wells at each dilution 21 days later. The other half of the culture was mechanically lysed and treated identically to quantify virus produced during the latency culture period (pre-reactivation) [42]. WT HCMV, but not the ΔmiR-US22 mutant, was able to reactivate in CD34+ HPCs (Fig 7). These data indicate that expression of miR-US22 during reactivation in CD34+ HPCs is necessary to produce infectious virus. These findings, together with the ability of miR-US22 to decrease EGR-1 expression resulting in altered proliferation and differentiation of CD34+ HPCs, indicate that the miRNA is necessary to alter cellular homeostasis in order to favor viral reactivation.

**Fig 7.**
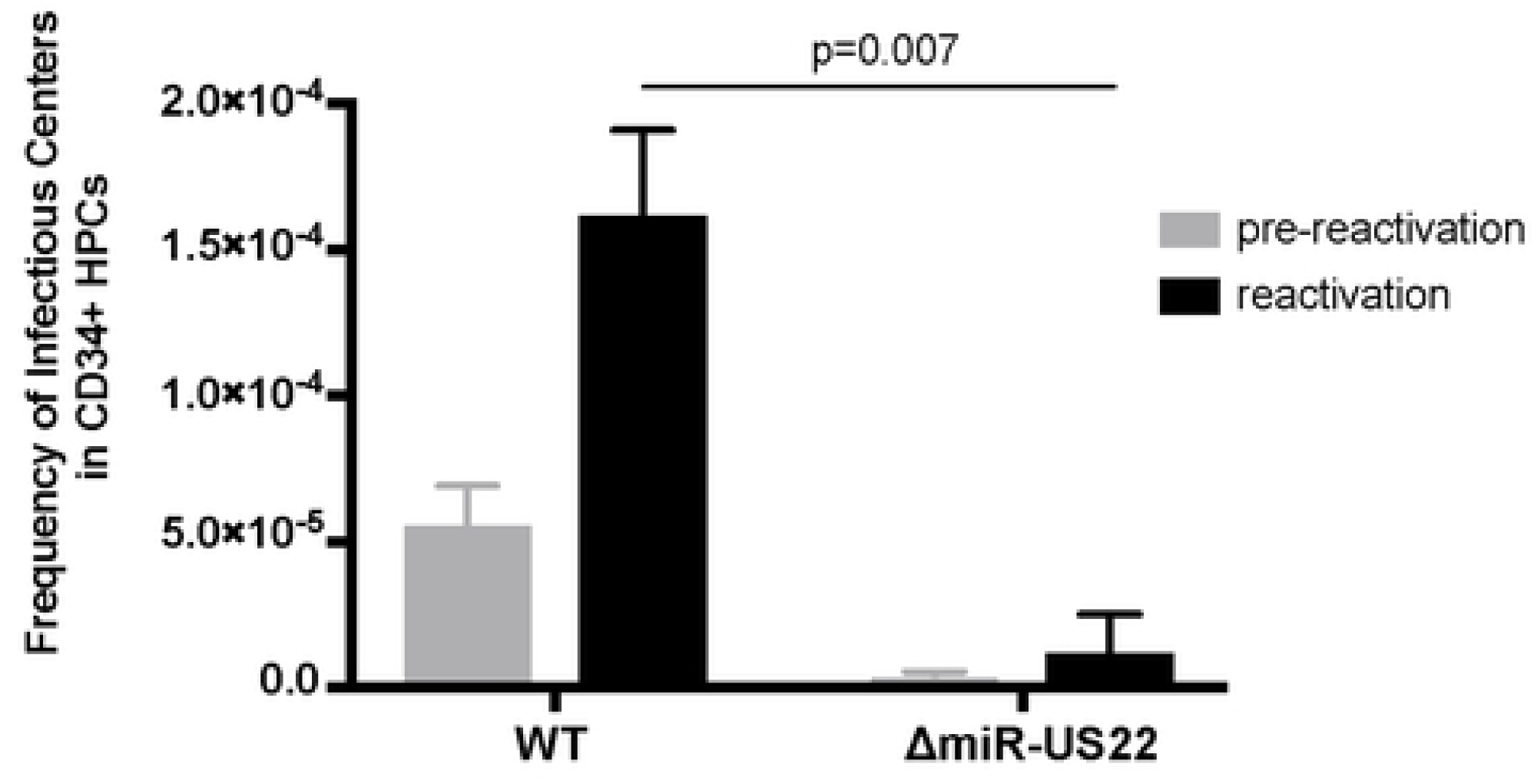
miR-US22 is required for viral reactivation in latently infected CD34+ HPCs. CD34+ HPCs were infected with WT TB40E or ΔmiR-US22 TB40E for 48 hours, and FACS-isolated for viable CD34+ GFP+ HPCs. Sorted cells were plated on stromal cell support for 12 days to establish viral latency. Viral reactivation was induced by co-culture on fibroblasts with cytokine stimulation for 21 days, and reactivation was measured by ELDA assay. The pre-reactivation control represents the amount of virus present in cell lysates at the end of latency prior to beginning reactivation. Reactivation data shown are from 2 independent experiments. Significance was calculated using paired t-test.

## Discussion

In this study we demonstrate that the transcription factor EGR-1 is directly targeted for down-regulation by HCMV miR-US22 that results in decreased proliferation of CD34+ HPCs and a decrease in total hematopoietic colony formation. We also show that an HCMV miR-US22 mutant fails to reactivate in CD34+ HPCs, indicating that expression of EGR-1 inhibits viral reactivation during latency. Since EGR-1 promotes CD34+ HPC self-renewal in the bone marrow niche, HCMV down-regulation of EGR-1 is a necessary step to block HPC proliferation and induce the cellular differentiation necessary to promote reactivation of virus. We propose a model of HCMV reactivation in CD34+ HPCs, in which latently infected cells initiate a process of reactivation that results in expression of miR-US22 (Fig 8). Subsequently, HCMV miR-US22 down-regulates expression of EGR-1 that results in the mobilization and differentiation of CD34+ HPCs from the bone marrow compartment to the peripheral blood to become monocytes that undergo further differentiation into macrophages, possibly due to expression of UL7 [43].

**Fig 8.**
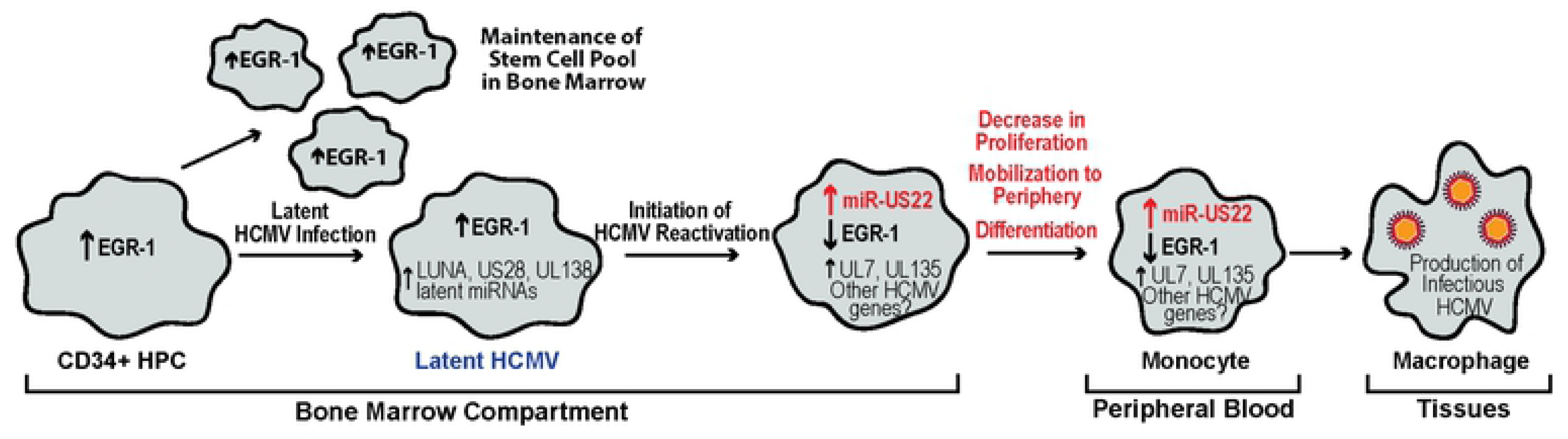
Model of the role of miR-US22 in HCMV reactivation. In the bone marrow, elevated EGR-1 levels in uninfected CD34+ HPCs contribute to the maintenance and self-renewal of these cells. In HCMV infected HPCs, stem cell quiescence and retention in the bone marrow contribute to the maintenance of the latency program. This stage is marked by the expression of known latency-promoting viral factors such as LUNA, US28, UL138, and miR-UL148D. Because HCMV miR-US22 expression is turned off, EGR-1 levels are high, which promotes viral latency by maintaining the undifferentiated state of the infected cell. As HCMV reactivation from latency is initiated, miR-US22 expression is induced, which results in decreased EGR-1 levels. Low EGR-1 levels induce mobilization of the infected cells - an event associated with viral reactivation.

HPCs are predominantly found in the bone marrow compartment that provides an environment in which the cells remain in a state of dormancy until hematopoietic stress induces cytokine signals that result in either cellular proliferation to replenish progenitors, or differentiation into myeloid or lymphoid cells to respond to infection. Total stem cell numbers are regulated by a balance between maintenance of quiescence and proliferation that is accomplished through apoptosis or migration of cells in and out of the bone marrow compartment [44, 45]. Transcription factors such as EGR-1, PU.1, runt related transcription factor 1 (RUNX1), kruppel-like factor 4 (KLF4), and CCAAT/enhance-binding protein alpha (C/EBPα) play key roles in the regulation of stem cell numbers in the bone marrow or differentiation into myeloid, lymphoid, or erythroid progenitor populations. Interestingly, cellular miRNAs are expressed in a cell type lineage specific manner to fine-tune expression of these transcription factors with both activation and feedback mechanisms (Gangaraju, 2009 #36265;[46]. Similarly, HCMV miR-US22 regulates EGR-1 levels in HPCs to inhibit proliferation and differentiation of the cell to create a cellular environment that promotes viral reactivation.

Expression of EGR-1 is finely tuned during normal hematopoiesis to directly regulate the hematopoietic processes of quiescence, apoptosis, proliferation, and differentiation. A number of published studies have, what at first glance appear to be contradicting roles for EGR-1 during these steps. However, since EGR-1 has very specific roles in different cell types and during different developmental stages, these data all contribute to a model in which EGR-1 specifically regulates and is regulated by distinct hematopoietic stages. In cultured cells, the overexpression of EGR-1 can both inhibit the differentiation of myeloid progenitors to the monocyte lineage [20] and induce terminal macrophage differentiation from the monocyte stage [21]. In CD34+ HPCs, Krishanaraju et al [20] show that overexpression of EGR-1 increases early stage differentiation of Blast-stage monocytes that correlates with the decrease in myeloid colony formation observed in Egr-1 knockout mouse bone marrow [47] and our results using knockdown of EGR-1 in human HPCs (Fig 5A). In vivo, Egr-1 -/- mice exhibit a significant decrease in progenitor cell proliferation in the bone marrow compartment and an increase in peripheral blood HPCs [48]. Therefore, EGR-1 plays an important role in stem cell quiescence, self-renewal, differentiation, and migration to the periphery. HPC quiescence and retention in the bone marrow niche are important events during HCMV latency. In this study, expression of HCMV miR-US22 blocks proliferation of HPCs due to down-regulation of EGR-1, which may allow for specific differentiation along the myeloid lineage [47], and provides a trigger for viral reactivation from latency. Therefore, high expression of EGR-1 in HPCs promotes viral latency by maintaining the undifferentiated state of the CD34+ HPCs. In contrast, low EGR-1 expression in infected HPCs induces mobilization and differentiation of progenitors to the myeloid lineage [47] that is associated with viral reactivation.

In addition to maintenance of the progenitor cells in the bone marrow niche, EGR-1 was recently shown to regulate expression of HCMV UL138 - a viral gene that is up regulated by EGFR signaling and is required to maintain the viral latent state (Buehler et al co-submission). Buehler et al observed that the UL138 promoter contains two EGR-1 binding sites, and mutation of one of these sites reduces UL138 expression. Consistent with a role for EGR-1 in regulating UL138 expression, co-transfection of a vector containing the UL133-UL138 region with miR-US22 resulted in reduced UL138 expression. Infection of fibroblasts with the HCMV ΔmiR US22 resulted in increased UL138 expression. These results indicate that miRUS22 regulates UL138 expression through EGR-1 and suggest that reduction of UL138 through miR-US22-mediated reduction of EGR-1 during reactivation from latency may be an additional step to reactivate virus.

Herpesvirus-encoded and cellular miRNAs have been shown to be important determinants for maintaining viral latency and reactivation. KSHV encodes miR-K9 that targets ORF50 (RTA) - the latent/lytic switch for the virus to maintain latency [49]. HCMV-encoded miR-UL112-3p was also shown to target the viral transcriptional activator IE72 (UL123) and UL112/113 that are necessary to activate early and late HCMV genes needed for viral reactivation [28]. Similarly, a cellular miRNA, neuron-specific miR-138, was shown to target HSV ICP0 that, when disrupted, allowed viral reactivation [37]. HCMV miR-UL148D was also reported to facilitate HCMV latency by inhibiting immediate early response gene 5 that promotes cell division cycle 25B protein and cyclin-dependent kinase 1-mediated suppression of IE72 [50]. EBV miRNAs were recently shown to regulate B Cell receptor signaling [51], and thus B cell activation, proliferation, and differentiation [52]. In this study, EBV miR-BHRF1-2-5p promotes latency by targeting GRB2 that is part of a signaling cascade that activates transcription factors, such as NFκB and Jun that induce genes that participate in B cell proliferation and survival. EBV-encoded miRs-BHRF1-2-5p and miR-BART2-5p, and a cellular miRNA miR-17-5p restrict lytic reactivation by dampening cellular responses to BCR cross-linking [52]. HCMV, similar to EBV, also regulates cellular differentiation in progenitor cells to maintain the viral latency state or to reactivate following differentiation. However, rather than regulating signaling pathways that activate transcription factors to reprogram the cellular differentiation program, HCMV directly targets EGR-1 - a central regulator of progenitor cell homeostasis.

Analysis of HCMV miRNA expression indicates that only subset of the 14 viral miRNAs (miRs -UL22A, -UL112-3p, -UL148D, -US5-1, -US5-2, -US25-1, and -US25-2-3p) are expressed in latently infected CD34+ HPCs. While the functions of these miRNAs in CD34+ HPCs during latency are unknown, the miRNAs most likely play important roles in maintenance of the virus during latency. HCMV miR-US22 is not expressed during latency in CD34+ HPCs but represents a class of viral gene products that are unnecessary for replication in fibroblasts but are required to initiate viral reactivation in CD34+ HPCs through the induction of cellular proliferation and differentiation. Another member of this class of HCMV genes is UL7, an early-late gene that functions as an Flt-3 receptor ligand [43]. Similar to miR-US22, deletion of UL7 does not alter viral replication in fibroblasts but blocks viral reactivation in CD34+ HPCs through induction of cellular differentiation. Therefore, HCMV encodes genes needed to reprogram the cell to allow the expression of cellular and viral genes necessary for viral replication. The identification of HCMV gene products like UL7 and miR-US22 that are necessary for viral reactivation from latency may provide important targets for early therapeutic HCMV intervention with new classes of drugs.

## Materials and methods

### Cells and media

HEK293T (CRL-11268; ATCC) and adult normal human dermal fibroblasts (NHDF) (CC-2511; Lonza) were cultured in Dulbecco’s modified Eagle’s medium (DMEM) supplemented with 10% heat-inactivated fetal bovine serum (FBS; HyClone), 100 units/ml penicillin, and 100ug/ml streptomycin (Invitrogen). Human aortic endothelial cells (AEC) (CC-2535; Lonza) were cultured in EBM-2 basal medium with EGM-2 SingleQuots^TM^ supplement excluding Heparin (Lonza), as well as 10% FBS, penicillin, and streptomycin. All cells were cultured at 37**°**C in 5% CO2.

### HCMV constructs

Wild type HCMV TB40/E-GFP bacterial artificial chromosome (BAC), in which the SV40-GFP cassette was introduced as a marker for infection was used to generate infectious virus [55, 56]. Mutant virus containing nucleotide changes in the miR-US22 stem loop of the WT TB40E BAC was constructed using galK recomineering. Briefly, the galactokinase (galK) gene was inserted within miR-US22-5p using homologous recombination, and was then replaced with the following annealed oligos to introduce the desired mutations that disrupt miR-US22 hairpin formation, but do not interfere with US22 ORF expression: GGTCTGGTCCGTCGTCTCCCATCTGGTCGGGTTCGGGGATGGGGAC CTC AAG CAA CGTGTGTCCGCGGGCGTGCATGGCTTTTGCTCGCCGGCCGCGCTG and CAGCGCGGCCGGCGAGCAAAAGCCATGCACGCCCGCGGACACACGTTGCTTGAGGTCCCCATCCCCGAACCCGACCAGATGGGAGACGACGGACCAGACC. All virus stocks were propagated and titered on NHDFs. For viral growth curves, NHDFs were infected at 3 pfu/cell for single step and 0.05 pfu/cell for multi step for 2 hours. Both cell-associated and supernatant viruses were harvested at multiple timepoints, and titers were determined by plaque assay on NHDFs. For all other infections, NHDF and AEC were inoculated with 3 pfu/cell for 2 hours at 37C. Afterwards, the viral inoculum was removed and replaced with fresh medium. Samples were harvested as indicated for each experiment.

### Reagents

The 3’UTR of human Egr-1 was amplified by PCR from genomic DNA extracted from NHDFs using DNAzol, and was cloned downstream of the *Renilla* luciferase gene in the pSICHECK2 plasmid (Promega) by XhoI and NotI restriction sites. Mutations in the seed sequence of the miR-US22 target site in Egr-1 were introduced by site-directed mutagenesis using the following primer pair: GAACTTGGACATGGCTGTTGGAGGCAGCTGAAGTCAAAGG and CCTTTGACTTCAGCTGCCTCCAACAGCCATGTCCAAGTTC. The mutated construct was verified by sequencing. Short-hairpin RNA (shRNA) targeting EGR-1 was cloned into pSiren expression plasmid via BamHI and EcoRI restriction sites using the following sequence: TGCTGTTGACAGTGAGCGATCCAGAATGTAAGAAAACAAATAGTGAAGCCACAGATGTATTTGTTTTCTTACATTCTGGAGTGCCTACTGCCTCGGA [57]. miR-US22 expression cassette was amplified from TB40E WT viral DNA and cloned into pSiren via BamHI and EcoRI restriction digest, using the following primer pair: GGCGGATCCCGGGGAAAGGGAATCTGCTTTTAG, and GGGAGAATTCGAAAACGAGGACGACACGAC. Cignal EGR-1 reporter kit (Qiagen) was transfected in HEK293T following manufacturer protocol. siGENOME RISC-Free control siRNA (NEG; Dharmacon) and EGR-1 siRNA (s4538; Thermofisher) were purchased for use in transfection experiments. Double stranded miRNA mimics were custom designed and synthesized by IDT (Integrated DNA technologies). PODS human EGF (Cell Guidance Systems) was dissolved in water for 100ug/mL stock solution and used at the indicated final concentrations for each experiment. The following commercial antibodies were used: EGR-1 (A303-390A-M, Bethyl Laboratories, Inc), GAPDH (ab8245, Abcam), CMV ICP22 for detection of the US22 gene product (sc-56974, Santa Cruz Biotechnology), and anti-CMV clone 8B1.2 for detection of IE72 (MAB810, MilliporeSigma™).

### Luciferase assays

HEK293T cells seeded into 96-well plates were co-transfected in triplicate with 100ng plasmid and 3pmol mimic per well using Lipofectamine 2000 (Invitrogen). Twenty hours later, the cells were harvested for luciferase assay with the Dual-Glo reporter assay kit (Promega). Luminescence was detected using Veritas microplate luminometer (Turner Biosystems). For EGR-1 reporter assays, 16 hours post transfection, cells were serum-deprived in 0%FBS DMEM for 4 hours, and then treated with 5 ng/mL EGF for 4 hours. All experiments were performed in triplicate and results are shown as mean ± standard deviation.

### Immunoblotting

Cells were harvested in protein lysis buffer (50mM Tris-HCl pH 8.0, 150mM NaCl, 1% NP-40, and protease inhibitors). Cell lysates, along with loading buffer (4X Laemmli Sample Buffer with 2-mercaptoeathanol) were incubated at 95C for 5 minutes, loaded on 4-20% polyacrylamide gels (BioRad), and transferred to Immobilon-P Transfer Membranes (Millipore Corp). After visualizing protein levels with the specified antibodies, the relative intensity of bands was quantitated using Fiji software (https://fiji.sc).

### Quantitative RT-PCR

Reverse transcription PCR (RT-PCR) was used to quantify viral microRNA expression in infected NHDF or CD34+ HPCs. Total RNA was isolated from infected cells using Trizol following the manufacturer’s instructions. cDNA was generated with MultiScribe™ reverse transcriptase (Thermofisher) using 100 ng total RNA and custom-designed miRNA hairpin-specific primers. Samples were incubated at 16°C (30min), 42°C (30 min), and 85°C (5 min). ABI StepOnePlus real time PCR machine was used with the following program: initial denaturation at 95°C (10 min), and 40 cycles at 95°C (15 sec), 60°C (1 min). The reaction was performed with Taqman Fast Advanced master mix (ABI). HCMV miRNA primers and probes were custom designed (using sequences from miRBase and (Stark, 2012 #4912)). Viral miRNA expression was normalized to cellular miR-16 levels (Assay ID 000391; ABI). For quantitation of miRNA copy number in infected CD34+ HPCs, purified oligos representing the mature form of each miRNA were included in an independent RT reaction in a known quantity. For qPCR, serial dilutions of the RT reaction were included to determine absolute miRNA copy number.

### Limiting dilution reactivation assay

CD34^+^ HPCs isolated from human bone marrow were infected at an MOI of 2 for 20h in IMDM supplemented with 10% BIT9500 serum substitute (Stem Cell Technologies, Canada), 2 mM L-Glutamine, 20 ng/ml low-density lipoproteins (Calbiochem), penicillin/streptomycin, and 50 µM 2-mercaptoethanol. Following infection, pure populations of infected CD34^+^ HPCs (>98% GFP-positive) were isolated by fluorescence-activated cell sorting (FACSAria, BD Biosciences Immunocytometry Systems, San Jose, CA) using a phycoerythrin-conjugated antibody specific to CD34 (BD Biosciences). Cells were sorted by the University of Arizona Shared Service at the University of Arizona Cancer Center. Pure population of infected HPCs were cultured in trans-wells above an irradiated (3000 rads, ^137^Cs gammacell-40 irradiator type B, Best Theratronics, Ottawa, Canada) M2-10B4 and Sl/Sl stromal cell monolayer [58] for 10-12 days in Myelocults (Stem Cell Technologies) containing 1 µM hydrocortisone and penicillin/streptomycin. The frequency of infectious centers production was measured using a limiting dilution assay as described previously [59]. Briefly, infected HPCs were serially diluted 2-fold in α-MEM with 20%FBS, 1 µM hydrocortisone, 0.2 mM i-inositol, 0.02 mM folic acid, 0.1 mM 2-mercaptoethanol, 2 mM L-glutamine, and penicillin/streptomycin supplemented with 15 ng/mL each of Interleukin-6, granulocyte colony stimulating factor, and granulocyte-macrophage colony stimulating factor (R&D Systems, MN). Aliquots of 0.05mL of each dilution were added to 12 wells (first dilution corresponds to 20,000 cells per well) of a 96-well tissue culture plates containing MRC-5 cells. To differentiate virus made as a result of reactivation from virus pre-existing in the long-term cultures, an equivalent number of cells were mechanically disrupted and seeded into MRC-5 co-cultures in parallel to the reactivation experiments. MRC-5 cells were monitored for GFP expression for a period of 14 days. The frequency of infectious centers formed was calculated based on the number of GFP^+^ cells at each dilution using software, Extreme limiting dilution analysis (ELDA, http://bioinf.wehi.edu.au/software/elda) [60]. For latency assays, CD34^+^ human hematopoietic cells (HPCs) were isolated from de-identified medical waste following bone marrow harvest from normal donors for clinical procedures at the University of Arizona Banner Medical Center.

### CD34+ HPC transfection, proliferation, and differentiation assays

Primary CD34+ HPCs were thawed and recovered overnight in stem cell media (IMDM containing 10% BIT serum replacement (Invitrogen), penicillin/streptomycin and stem cell cytokines (SCF, FLT3L, IL-3, IL-6). Following recovery, HPCs were transfected using the Amaxa 4D system and the Primary Cell P3 solution according to the manufacturer’s instructions (Lonza). HPCs were transfected with 1ug pSIREN plasmid DNA per 10^6^ cells using either program EH-100 or EO-100. HPCs were recovered in stem cell media for 48hrs, then isolated by FACS (BD FACS Aria equipped with 488, 633 and 405 lasers, run FACS DIVA software) for a pure population of viable, CD34+, GFP+ HPCs. Pure populations of sorted HPCs were plated either at 500 cells/mL in Methocult H4434 (Stem Cell Technologies) in 6 well plates in triplicate for myeloid colony assays, or at 10^4^ cells/mL in stem cell media, 200uL/well in 96 well plates for proliferation assays. Myeloid colonies were enumerated at 7 and 14 days using a standard microscope. Total and specific colony types were determined manually. Proliferation was assessed at indicated times by Trypan Blue exclusion and manual counting.

### Statistical analysis

Data are shown as mean ± standard deviation. Statistical analysis was performed using GraphPad Prism (v6 or v7) for comparisons between experimental groups using unpaired t test or two-way analysis of variance (ANOVA) with Tukey’s post-hoc test.

### Ethics Statement

CD34+ hematopoietic progenitor cells (HPCs) were isolated from de-identified human fetal liver obtained from Advanced Bioscience Resources as previously described [53] or were isolated from de-identified medical waste following bone marrow isolations from healthy donors for clinical procedures at the Banner-University Medical Center at the University of Arizona as previously described [54].

## Acknowledgements

This work was supported by grant AI21640 and P01AI127335 (JAN), R01 AI079059 and P01AI127335 (FG) from the National Institute of Allergy and Infectious Diseases, and an American Cancer Society Post-Doctoral Research Fellowship (129842-PF-16-212-01-TBE) funded to J.B. We are grateful for helpful discussions with Patrizia Caposio, Dan Streblow, and Andrew Yurochko.

